# Tissue factor-dependent colitogenic CD4+ T cell thrombogenicity is regulated by activated protein C signalling

**DOI:** 10.1101/2024.04.16.589774

**Authors:** Gemma Leon, Paula A. Klavina, Aisling M. Rehill, Shrikanth Chomanahalli Basavarajappa, James S. O’Donnell, Seamus Hussey, Patrick T. Walsh, Roger J.S. Preston

**Affiliations:** Irish Centre for Vascular Biology, School of Pharmacy and Biomolecular Sciences, RCSI University of Medicine and Health Sciences; National Children’s Research Centre, Crumlin, Dublin; Department of Paediatrics, RCSI University of Medicine and Health Sciences, Dublin; Department of Clinical Medicine, Trinity Translational Medicine Institute, Trinity College Dublin

## Abstract

Inflammatory bowel disease (IBD) patients experience up to 6-fold increased risk of venous thromboembolism (VTE) compared to the general population, although the mechanistic basis for this increased risk remains poorly defined. We found that colitogenic CD4^+^ T cells express tissue factor (TF) and promote rapid TF-dependent plasma thrombin generation in T cell-dependent calibrated automated thrombinography assays. Furthermore, we identified the presence of TF^+^CD4^+^CD3^+^ T cells in the colons of both mice with colitis and paediatric IBD patients during active disease. TF is typically expressed in an ‘encrypted’ state and requires decryption for optimal procoagulant activity. Notably, flow cytometric analysis demonstrated that activated CD4^+^ T cells express significantly increased acid sphingomyelinase and protein disulphide isomerase, critical mediators for TF decryption, on their cell membrane compared to naïve T cells. The protein C (PC) pathway is an important regulator of TF-mediated thrombin generation. Pertinently, pre-clinical studies suggest an important role for diminished PC pathway activity in IBD pathophysiology. To understand how this process might be regulated, we performed meta-transcriptomic and gene expression analysis of IBD patient gut biopsy tissue, identifying dysregulated expression of genes involved in the regulation of coagulation, including PC (*PROC)* and its receptor (EPCR; *PROCR)*. Subsequent functional studies revealed that activated protein C (APC) signalling reduced colitogenic T cell generation and activity, potently impaired TF decryption and significantly reduced T cell-mediated thrombin generation and clot formation. These data identify TF-mediated colitogenic T cell thrombogenicity and demonstrate a new role for APC signalling in regulating T cell thrombo-inflammatory activity.

## BACKGROUND

Inflammatory bowel disease (IBD) is a chronic inflammatory disorder of the gastrointestinal (GI) tract, commonly sub-grouped into Crohn’s Disease (CD) and Ulcerative Colitis (UC)^1^. Approximately 10 million people suffer from this debilitating disease worldwide, and the incidence is rising^2^. The pathology of IBD, particularly CD, is largely mediated by aberrant T cell responses^1^. Despite the emergence of biologics targeting T cell proinflammatory activity, many patients remain unresponsive to these therapies, and many become resistant over time^3^. IBD patients are at greater risk of developing venous thromboembolism (VTE) compared to the general population. Adults with IBD exhibit a 3-fold higher risk of VTE^4^, and this risk rises to 6-fold higher in paediatric IBD patients^4,5^. Strikingly, 22% of IBD patients who do develop VTE, die within 2 years of the first event^6^, and 33% experience recurrent VTE within 5 years^7^. Several factors may contribute to this increased risk, including disease severity, microbial dysbiosis, pregnancy, treatment and GI surgery^8,9^.

Despite the explicit association between VTE and IBD, the molecular pathogenesis of VTE in IBD patients remains poorly understood. Plasma levels of several acute phase procoagulant and anti-fibrinolytic proteins, including von Willebrand factor, plasminogen activator inhibitor 1 and fibrinogen, are elevated during IBD ‘flares’^10^, whereas anticoagulant proteins such as protein C (PC), protein S and antithrombin are diminished^11^. In addition, leukocytes and platelets from IBD patients have been reported to shed increased numbers of procoagulant tissue factor (TF)^+^ microparticles^12,13^. Notably, antibody-mediated TF inhibition decreases disease activity in mice during dextran sodium sulphate (DSS)-induced colitis, with reduced leukocyte and platelet adhesion to colonic venules, and significantly lower rates of thrombus formation^11^. Dysregulation of the anticoagulant and anti-inflammatory PC pathway has also been implicated in IBD pathogenesis. PC^low^ mice develop spontaneous colitis^14^, and mice largely deficient in endothelial protein C receptor (EPCR), display exacerbated disease in pre-clinical IBD models^15^. This has been postulated to arise due to decreased EPCR and thrombomodulin (TM) expression in the inflamed microvasculature of mice with colitis, resulting in a reduced capacity for PC activation^14,16^.

Colitogenic CD4^+^ T cell responses are critical for the induction and perpetuation of IBD^17^. Although largely overlooked in the context of thrombosis, recent studies have indicated a role for T cells in thrombus development and regulation. For example, a novel Treg subpopulation that produces secreted protein acidic and rich in cysteine (SPARC) was found to promote thrombolysis via recruitment of CD11c^+^ monocytes with enhanced fibrinolytic activity to the thrombus^18^. Conversely, T cell-targeted immunotherapies, such as immune checkpoint inhibitors, have been reported to confer an increased risk^19^ and incidence^20^ of VTE in cancer patients, although the mechanisms underlying this phenomenon are currently unknown.

In this study, we demonstrate a novel role for CD4^+^ T cells in facilitating thrombin generation and clot formation via T cell activation-induced TF activity. We show that TF expression is upregulated in the intestinal tissue of IBD patients and mice with colitis, and report for the first time the elevated presence of CD4^+^TF^+^ T cells in the gut mucosa during colitis. Furthermore, we demonstrate APC cell signalling mitigates pro-thrombotic T cell responses, independent of its canonical anticoagulant activity. These data implicate enhanced T cell-mediated thrombogenicity as a potential mediator of increased VTE risk in IBD patients and highlight a new role for APC anticoagulant signalling in regulating colitogenic T cell thrombo-inflammatory activity.

## MATERIALS AND METHODS

Further detailed methodology can be found in the ‘*Supplemental Methods and Materials’*.

### Study subjects

All human samples were obtained with consent/assent from paediatric IBD patients and control participants recruited in the <u>D</u>eterminants and <u>O</u>utcomes of <u>CH</u>ildren and <u>A</u>dolescents with IBD <u>S</u>tudy (DOCHAS) at the gastroenterology unit at Children’s Health Ireland (CHI), Crumlin (Dublin, Ireland). All participants underwent diagnostic evaluation according to international paediatric standards (Porto criteria) and rigorously phenotyped using the paediatric-specific Paris classification of IBD. Rectal and colonic biopsies were obtained from patients enrolled in the study. Patients initially enrolled with suspected IBD, but subsequently not diagnosed with disease, comprise the control population. All experiments using these tissues were performed under approval from the institutional Research Ethics Committee (GEN/193/11), and included 30 participants (CD, n=11; UC, n = 7; Ctrl, n = 12).

### Isolation and culture of CD4^+^ T cells

Anonymised healthy donor buffy coats were obtained from the Irish Blood Transfusion Service, St. James’ Hospital, Dublin. PBMCs were isolated using Lymphoprep density gradient centrifugation (STEMCELL Technologies), and negative magnetic selection was then used to purify CD4^+^ T cells (CD4 T cell isolation Kit, human, Miltenyi Biotec). Isolated CD4^+^ T cells were plated at a density of 0.8x10^6^/ml in AIM media supplemented with CTS Immune Cell SR (Gibco, ThermoFisher), activated with anti-CD3/anti-CD28 activation beads per manufacturer’s instructions (Gibco Dynabeads Human T-Activator CD3/CD28 for T Cell Expansion and Activation, Thermofisher), stimulated with IL-2 (R&D), and/or T cell differentiating cytokines and antibodies (Th1: anti-IL-4, IL-12 (Miltenyi Biotec), Treg: TGFβ (Immunotools), Th17: TGFβ, IL-1β (R&D), IL-23 (R&D), IL-6 (R&D), anti-IFNγ (Miltenyi Biotec)) +/-APC (Cambridge ProteinWorks) and then incubated at 37 °C for 5-7 days.

### T cell transfer mediated colitis model

T effector cells (CD4^+^ CD25 ^−^ CD45RB^hi^) were FACs sorted from C57BL/6 mice and injected *i.p*. into *Rag1^−/−^* recipient mice (5 × 10^5^). Littermate control *Rag1^−/−^* recipient mice were injected *i.p*. with PBS. Disease progression was measured by percentage weight loss compared to the original weight, with a 20% loss of original weight at the cut-off point. Colons were harvested for analysis when clinical signs of colitis were evident (4 weeks post transfer), and histology was performed to confirm colitis. Harvested colon tissue was fixed in 10% formalin overnight, dehydrated and embedded into paraffin blocks. Paraffin blocks were sectioned to 5 μm thickness using a microtome, mounted on Superfrost Plus adhesion slides (Thermofisher), and stained with haematoxylin and eosin (Thermofisher). Histological disease scoring was performed blinded, as described previously^21^.

All mice were housed under specific pathogen-free conditions, on a 12-hour light/dark cycle, in a temperature-controlled unit at the Comparative Medicine Unit in Trinity Translational Medicine Institute, St. James Hospital, Dublin, Ireland. Food and water were provided *ad libitum*. The Health Products Regulatory Authority approved all animal experiments under Project License AE19136/P125.

### CD4^+^ T cell-based thrombin generation assay

Following T cell culture, supernatants were removed, and cells were washed three times with PBS containing EDTA. 20µL MP-reagent (Stago) and 80µL of human normal pooled platelet-poor plasma (NPP) (Fanin), factor XII (FXII)-deficient plasma (HTI), or factor VII (FVII)-deficient plasma (HTI) were added to the washed T cells. The assay was then initiated with 20µL of FluCa (Stago), and fluorescence was recorded using Thrombinoscope software.

### CD4^+^ T cell-based clot formation assay

Following T cell culture, cell supernatants were removed, and cells were washed three times with PBS containing EDTA. 80µl of normal pooled plasma (38%), TBS-T (52%) and phospholipids (16µM) (10%) were added to the cells. The assay was then initiated with 20µL of CaCl_2_ (10.6mM; Sigma) and was read at OD_405_ and recorded in a kinetic assay every minute for 1 hour using a Synergy MX microplate reader (BioTek).

### Factor Xa generation assay

T cell surface TF activity was measured by T cell-mediated factor Xa (FXa) generation. Following T cell culture, cell supernatants were removed, and cells were washed three times with Buffer A (10mM Hepes, 0.15 M NaCl, 4 mM KCl, 11 mM glucose, pH 7.5). Cells were then incubated at 37 °C with 200ul of Buffer B (Buffer A, 5mM CaCl_2_, 1mg/ml BSA) containing FVIIa (10nM; Prolytix, Cambridge Biosciences) and FX (175nM; Cambridge Proteinworks) for 30 minutes. 25µl of the supernatant _w_as removed and added to 50µl TBS:BSA buffer (50mM Tris-HCL, 0.15 mol/L NaCl, 1mg/ml BSA, 25mol/L EDTA, pH 7.5). 50ml of this mixture was transferred to a 96-well plate, and 50µl of Factor Xa Chromogenic Substrate (BIOPHEN™ CS-11(65), Quadratech) was added. The rate of colour development was measured against known FXa concentrations (Cambridge Proteinworks) using a Synergy MX microplate reader (BioTek).

### Statistical analysis

For each dataset, statistical analysis was performed by first analysing the data for normal distribution (Shapiro–Wilk or Kolmogorov–Smirnov test) and equality of variance (F test). Unpaired data was then analysed by two-tailed Student’s t-test or one-way ANOVA for parametric data or Mann–Whitney U test for nonparametric data, as appropriate. Paired data was analysed by paired two-tailed student’s t-test for parametric data, or Wilcoxon matched-pairs signed ranks test for nonparametric data, as appropriate. All analysis and graph representation were performed using GraphPad Prism 9.5 software. Data are shown as means ± SEM.

## RESULTS

### The colonic tissue of IBD patients and mice with colitis is enriched with CD4^+^ TF^+^ T cells

To evaluate coagulation parameters in IBD patients, we examined publicly available transcriptomic data of colonic biopsies from paediatric IBD patients in the Risk Stratification and Identification of Immunogenetic and Microbial Markers of Rapid Disease Progression in Children with Crohn’s Disease (RISK) study^22^. We found that colonic tissue *F3* expression was significantly upregulated in CD patients, with either ileal (iCD) or colonic (cCD) sites of disease, when compared to non-IBD control participants (**Figure 1a**). Similarly, dysregulation in *F10, F2R, THBD, PROC, PROCR*, and *PROS1* gene expression was also observed **(Supplementary Figure 1a-e)**. Subsequent RNA-seq analysis of rectal biopsies from the DOCHAS paediatric IBD cohort during active disease revealed similar dysregulation in *F3* and other coagulation-associated genes in UC and CD patients compared to non-IBD patient biopsies (**Figure 1b**). KEGG pathway analysis suggested dysregulation in coagulation and complement pathways in the rectal tissue of CD and UC patients compared to the control population **(Supplementary Figure 2a and b).** Further *in silico* analysis of TF expression in human immune cells that contribute to IBD pathogenesis showed TF expression was markedly upregulated in CD4^+^ αβ T cells **(Supplementary Figure 3)**, leading us to question whether TF expression was present on CD4^+^ T cells in colonic biopsies from paediatric IBD patients. Notably, we found that CD4^+^TF^+^ T cells were significantly increased in the colonic tissue of IBD patients compared to patients who presented with GI discomfort, but were later shown not to have IBD (**Figure 1c and 1d**). These studies demonstrate that CD4^+^ TF^+^ T cells exist and are elevated in the gut of IBD patients.

**Figure 1:**
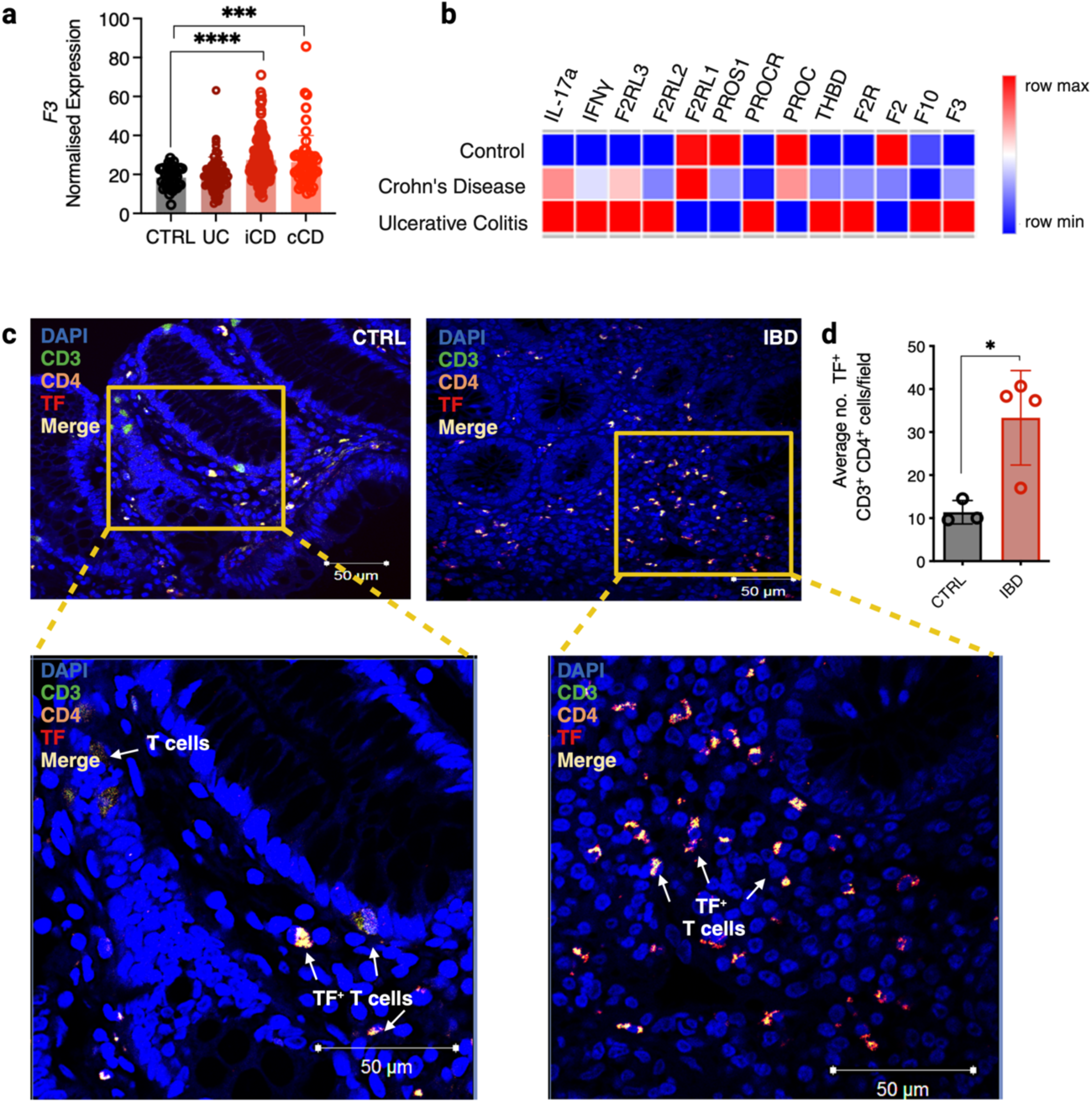
TF expression is upregulated during IBD. **(a)** Expression of TF from RNA-seq data of colonic biopsies from Crohns Disease (CD; illeal (iCD, n=162), colonic (cCD, n=56), Ulcerative Colitis (UC, n=62), and query IBD healthy control cohort (CTRL, n=42) patients in the RISK trial (GEO ID GSE57945). **(b)** Expression of coagulation and inflammation-associated genes from RNA sequencing of paediatric CD (n=11), UC (n=7), and query IBD healthy control colonic biopsies (n=12). **(c)** TF, CD3 and CD4 co-staining and **(d)** the average number of TF^+^ CD3^+^ CD4^+^ T cells per field in colonic biopsies from paediatric IBD patients (IBD, n = 4, non-IBD n=3). Student t-test or Mann Whitney U Test was used where appropriate to determine statistical significance with *P≤0.05, **P≤0.01, ***P≤0.001.

To investigate the potential role of TF in colitogenic T cells, CD4^+^CD25^low^CD45Rβ^high^ T effector cells were FACs sorted from C57B6 wild-type donor mouse splenocytes and injected into the peritoneum of *Rag1^-/-^* host mice. Disease progression was measured by weight loss. Once mice had lost 20% of their original body weight, the experiment was terminated, and colons were harvested for analysis (**Figure 2a**). After 4 weeks, clinical signs of disease were present in the T cell recipient group, as evidenced by the loss of 20% body weight (**Figure 2b**). H&E histology of colonic sections demonstrated that T cell-recipient mice displayed significantly increased disease scores compared to PBS recipients (p<0.0001; **Figure 2c**). Notably, TF expression was significantly upregulated in the colons of T cell transfer-recipient mice compared to vehicle-treated control mice (**Figure 2d-e**) and was specifically detected both in the inflamed colonic epithelium and on infiltrating cells. To evaluate whether infiltrating T cells were one of the cell subsets expressing TF in this tissue, we co-stained colon sections with CD3, the T cell receptor (TCR), and identified the significantly increased presence of TF^+^CD3^+^ T cells in the colons of mice during active colitis (**Figure 2f**).

**Figure 2:**
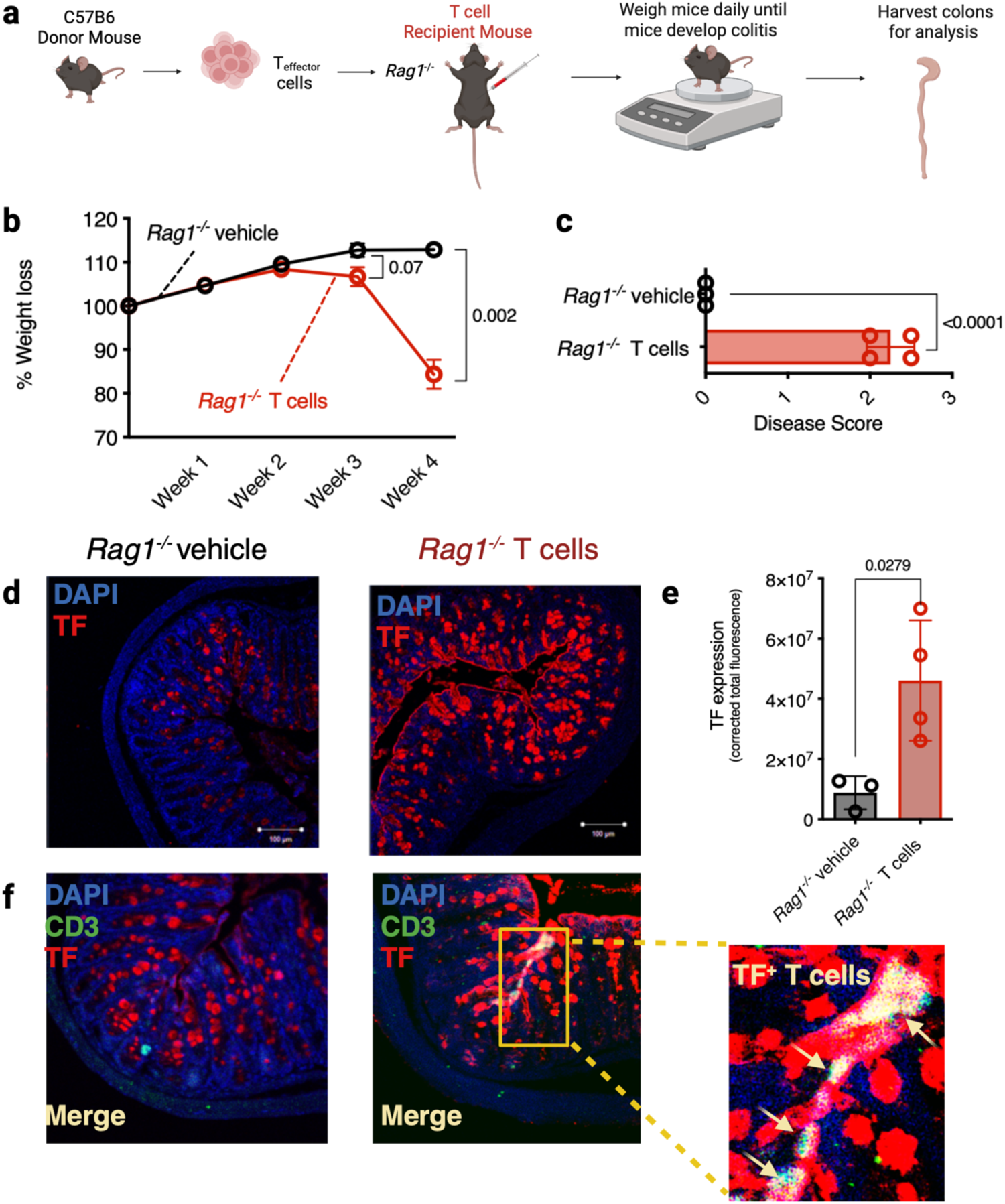
TF expression is enhanced in the colon and on infiltrating T_effector_ cells during T cell transfer-induced colitis. **(a)** Schematic diagram of the T cell transfer model of colitis. CD4^+^ T effector cells were isolated from donor C57BL/6 wild type (WT) mice and i.p. injected into host *Rag1^-/-^* mice. *Rag1^-/-^* mice *i.p.* injected with PBS were used as a control. **(b)** Colitis developed over a 4-week period, and disease progression was measured by % weight loss compared to original weight. **(c)** This was confirmed by subsequent colon histology analysis, **(d)** TF staining and **(e)** Corrected total fluorescence (CTF) of mice colons following T cell transfer-induced colitis. **(f)** TF and CD3 co-staining of mouse colons following T cell transfer-induced colitis. 2-way ANOVA or Student t-test was used where appropriate to determine statistical significance with *P≤0.05, **P≤0.01, ***P≤0.001.

### Activated inflammatory CD4^+^ T cells exhibit TF-mediated thrombogenic activity

To investigate the potential thrombogenic activity of TF^+^CD4^+^ T cells, we performed a bespoke CD4^+^ T cell-mediated thrombin generation assay that was designed so that cell-associated TF would be the sole trigger for thrombin generation (**Figure 3a**). We assessed isolated human unactivated naïve T cells (θ), TCR-activated T helper cells (Th0), and colitis-associated pro-inflammatory type 1 T helper cells (Th1) **(Supplementary Figure 4a)**. Interestingly, the presence of Th0 and Th1 cells promoted thrombin generation significantly more effectively than θ T cells, which had little effect (**Figure 3a-d**). Specifically, θ T cells exhibited a significantly longer lag time (**Figure 3b**), lower peak thrombin (**Figure 3c**), and a reduced endogenous thrombin potential (ETP; **Figure 3d**) compared to Th0 and Th1 cells. Similarly, using a plasma clotting assay, we also observed a significantly increased rate of plasma clot formation in the presence of Th0 and Th1 cells compared to naïve θ T cells (**Figure 3e**). To confirm that the increased rate and size of thrombin generation arose from T cell TF activity, the assay was repeated in the presence of FVII-deficient plasma, which prevented rapid Th0 and Th1-enhanced thrombin generation and therefore excluded a role for FXII activation in the observed cell-dependent thrombin generation (**Figure 3f-i**). In support of this, substitution with FXII-deficient plasma led to qualitatively similar results as when normal plasma was used **(Supplementary Figure 4b-e)**. Given the apparent role of activated T cell TF in enhanced thrombin generation, we assessed *F3* expression and TF surface expression in each T cell subset. *F3* expression (**Figure 3j**), TF protein expression (**Figure 3k-l**), and FXa generation (**Figure 3m**) were significantly increased on Th0 and Th1 cells compared to θ T cells. These data indicate that TF expression is upregulated in activated and colitogenic inflammatory T cells, facilitating enhanced extrinsic tenase activity and thrombin generation.

**Figure 3:**
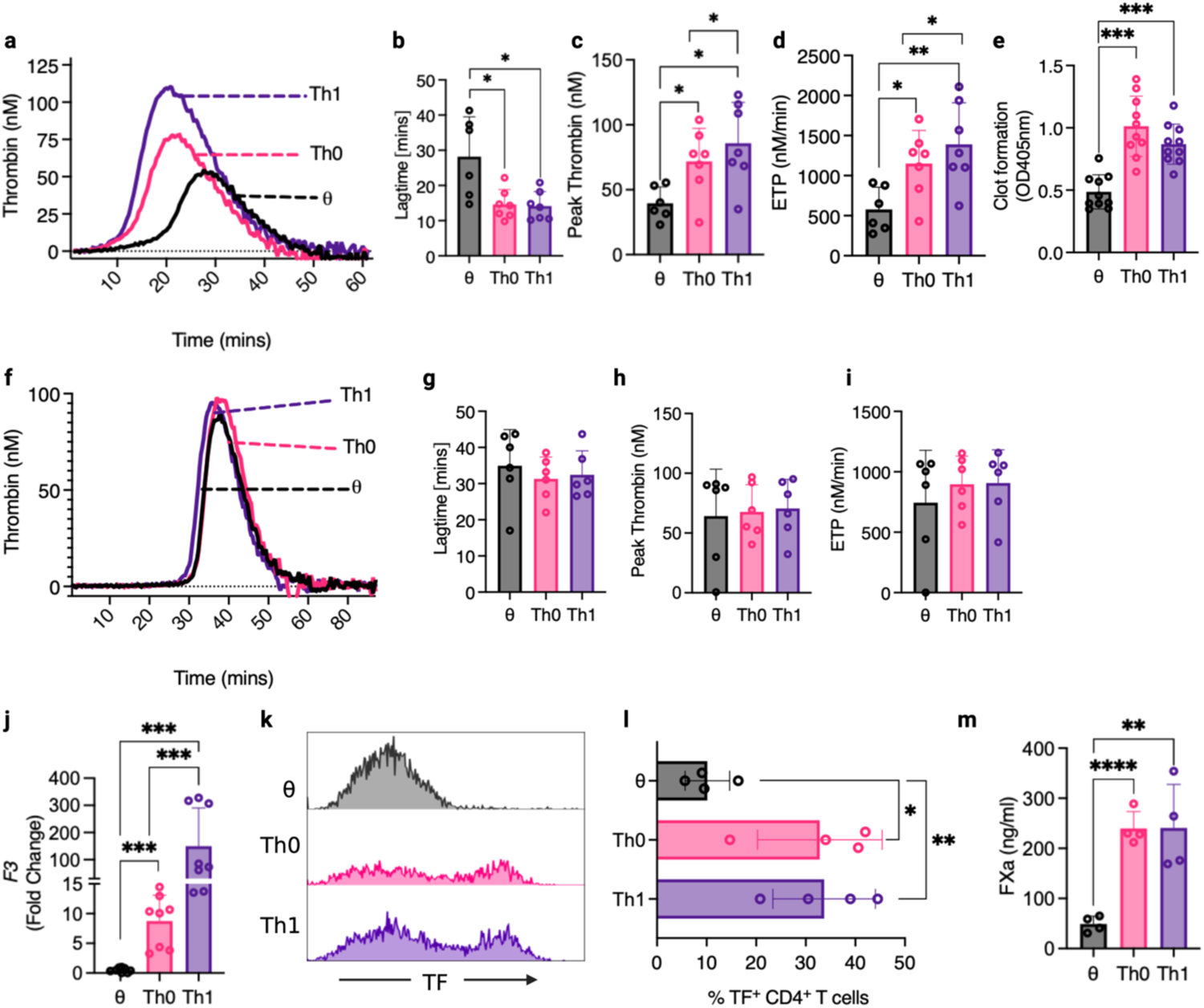
T cell activation enhances TF-mediated CD4^+^ T cell thrombogenicity. CD4^+^ T cells were isolated from donor human blood and plated at a density of 0.8x10^6^/ml with IL-2 for unactivated conditions (θ). Plated cells were activated by anti-CD3/anti-CD28 activation beads and stimulated with IL-2 for Th0 conditions, and differentiation cytokines (αIL-4 + IL-12) were added to skew cells to a Th1 lineage**. (a-d)** Following 5 days in culture, θ, Th0 and Th1 cells were washed with EDTA-containing PBS, and their ability to initiate thrombin generation was analysed by calibrated automated thrombinography in normal pooled platelet-poor plasma. **(b)** Lagtime, **(c)** peak thrombin levels and **(d)** endogenous thrombin potential (ETP) were measured and compared between θ, Th0 and Th1 cells**. (e)** The rate of clot formation was measured in θ, Th0 and Th1 cells. **(f-i)** θ, Th0 and Th1 cell-mediated thrombin generation was analysed by calibrated automated thrombinography using FVII-deficient platelet-poor plasma. **(g)** Lagtime, **(h)** peak thrombin levels and **(i)** ETP were measured and compared between θ, Th0 and Th1 cells. **(j)** *F3* gene expression, **(k&l)** cell surface TF expression and **(m)** FXa generation were measured in θ, Th0 and Th1 cells and compared. Student-paired t-test, Wilcoxon test, or Mann Whitney U Test was used where appropriate to determine statistical significance with *P≤0.05, **P≤0.01, ***P≤0.001 for 4-10 donors/group.

### CD4^+^ T cell activation promotes cell surface TF decryption

To investigate the mechanistic basis for T cell-mediated TF procoagulant activity, we evaluated molecular processes associated with enhanced TF activity, or ‘decryption’ in T cells. In the steady state, most immune cell surface TF is ‘encrypted’, and only adopts a procoagulant conformation in response to injury, inflammation, or cellular activation^23^. For example, the presence of the membrane phospholipid sphingomyelin (SM) in the outer leaflet of resting cells inhibits TF procoagulant activity^23^. However, the recruitment of acid-sphingomyelinase (ASMase) from lysosomes to the cell surface results in the hydrolysis of SM to ceramide and facilitates TF decryption (**Figure 4a**)^23^. Following T cell activation, we observed significantly increased ASMase trafficking to the cell surface of Th0 and Th1 cells compared to θ T cells (**Figure 4b-c**). Similarly, phosphatidylserine (PS) exposure on the outer leaflet increases TF procoagulant activity^23^ (**Figure 4d**). We therefore measured the binding of fluorescently-labelled PS-binding lactadherin to the surface of CD4^+^ T cells and found significantly increased lactadherin binding on Th0 and Th1 cells, compared to θ T cells (**Figure 4e-f**). Translocation of the oxidoreductase enzyme protein disulphide isomerase (PDI) to the cell surface contributes to TF decryption and procoagulant activity via disulphide bond formation and thiol exchange^23^ (**Figure 4g**). We found that T cell activation induces a significant increase in PDI expression on the cell surface of Th0 and Th1 cells compared to θ T cells (**Figure 4h-i**). Collectively, these data demonstrate that TF decryption mechanisms are responsive to T cell activation and support activation-induced TF expression and subsequent decryption as the mechanistic basis for CD4^+^ T cell-mediated procoagulant activity.

**Figure 4:**
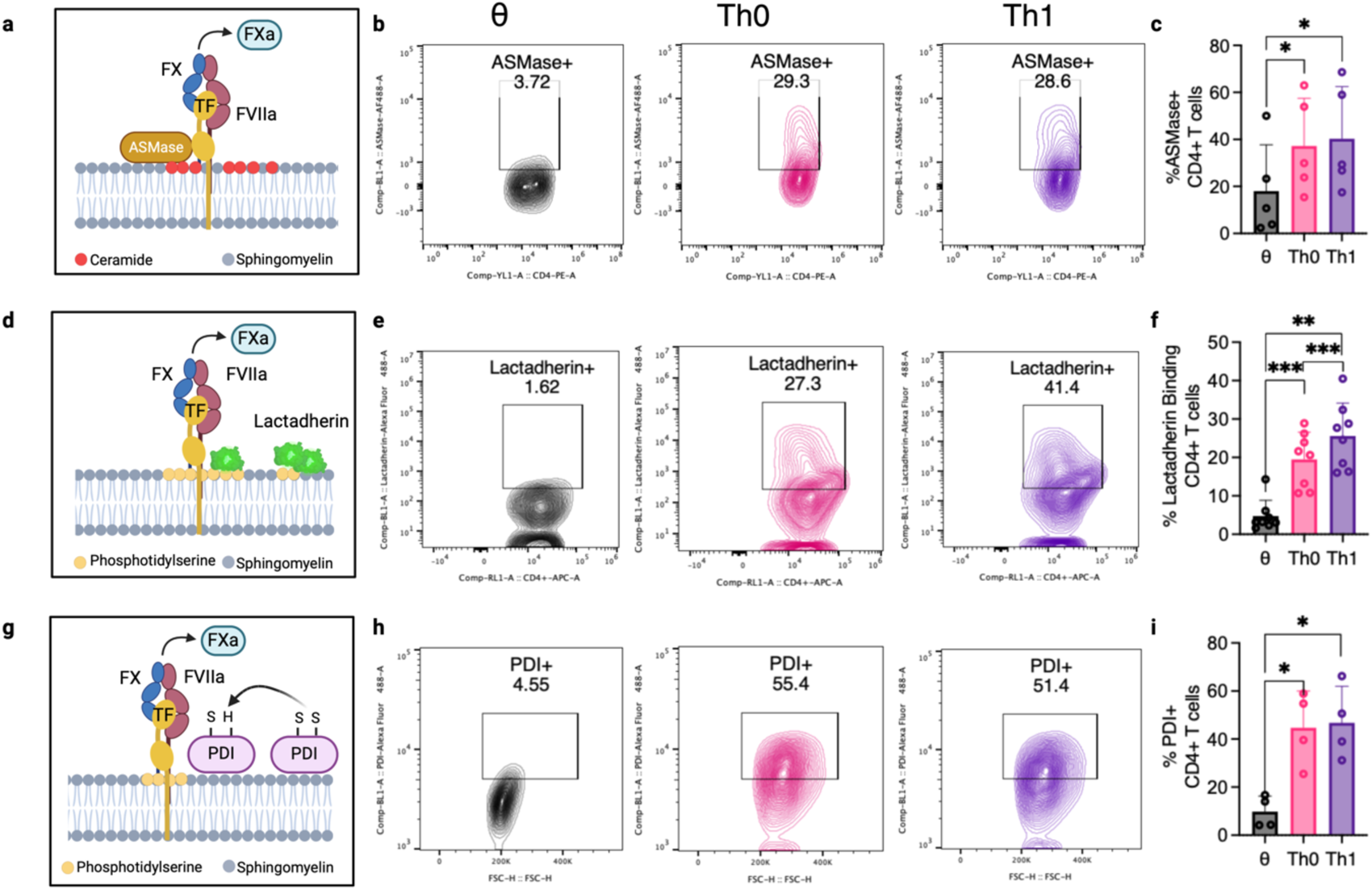
CD4^+^ T cell activation upregulates TF decryption. CD4^+^ T cells were isolated from donor human blood and skewed to θ, Th0, and Th1 cell subtypes as described previously. **(a)** ASMase translocation to the cell surface, **(d)** PS exposure on the outer membrane leaflet, and **(g)** PDI recruitment to the cell surface were assessed. TF decryption pathways in θ, Th0 and Th1 were analysed by **(b & c)** ASMase cell surface expression, **(e & f)** fluorescently labelled lactadherin binding to exposed cell surface PS, and **(h & i)** cell surface PDI expression. Student-paired t-test was used to determine statistical significance with *P≤0.05, **P≤0.01, ***P≤0.001 for 4-8 donors/group.

### The protein C pathway is dysregulated in IBD and regulates CD4^+^ T cell procoagulant activity

APC is an important regulator of TF-mediated thrombin generation^24^, IBD pathophysiology^14–16^, and T cell inflammatory responses^25–30^. Therefore we next sought to investigate a potential role for APC anti-inflammatory signalling in regulating TF^+^CD4^+^ T cell procoagulant activity. In addition to enhanced TF expression and activity, PC (*PROC*) gene expression was significantly reduced in the colonic biopsy tissue of IBD patients compared to a non-IBD patient cohort (**Figure 1b & Figure 5a**). APC bound to activated (Th0) and inflammatory (Th1) CD4^+^ T cells significantly better than θ T cells (**Figure 5b-c**) and, in keeping with prior studies, inhibited proinflammatory Th1 and Th17 differentiation whilst promoting the expansion of anti-inflammatory Tregs **(Supplementary Figure 5a-h).** To evaluate a potential regulatory role for APC, we treated each T cell subset with APC, washed the cells to remove residual APC binding, and assessed the capacity of APC-treated CD4^+^ cells to mediate T cell-mediated thrombin generation. Remarkably, APC pre-treatment of Th0 and Th1 cells also significantly decreased their capacity to facilitate thrombin generation (**Figure 5d-g, i-l)**. APC pre-treated Th0 or Th1 cells exhibited markedly longer lag time than untreated Th0 or Th1 cells (**Figure 5e & j)**, significantly lower peak thrombin (**Figure 5f & k)**, and significantly reduced ETP (**Figure 3g & l)**. Similarly, APC pre-treatment caused a significantly decreased rate of plasma clot formation in Th0 and Th1 cells (**Figure 5h & m)**. To confirm this APC-mediated anticoagulant effect was mediated by direct reduction of TF activity, we measured FXa generation on the surface of each T cell subset in response to APC pre-treatment. We observed a significant reduction in FXa generation on APC-treated Th0 and Th1 cells, compared to untreated T cells (**Figure 5n & r)**. To explore this further, we analysed TF expression in APC pre-treated CD4^+^ T cells. We found that TF (*F3*) expression was significantly reduced in both Th0 and Th1 cells treated with APC (**Figure 5o & s)**, and that these cells also displayed significantly lower levels of PDI on their surface (**Figure 5p-q, t-u)**. Taken together, these data demonstrate a novel non-canonical mechanism of APC anti-immunothrombotic activity, namely the direct inhibition of TF-mediated CD4^+^ T cell thrombo-inflammatory activity.

**Figure 5:**
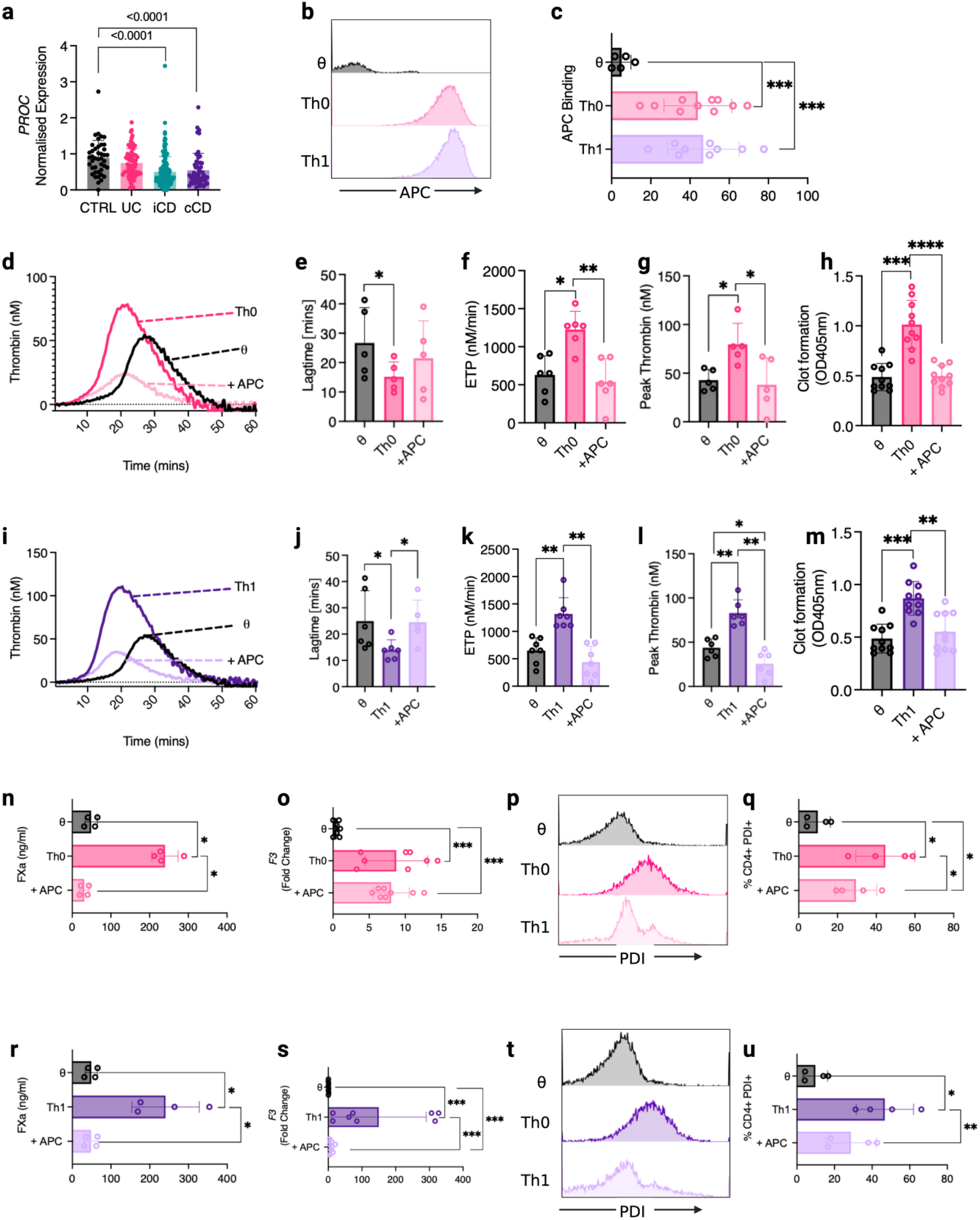
APC is downregulated in IBD and limits thrombogenic CD4^+^ T cell activity. **(a)** *PROC* (Protein C; PC) expression in colonic biopsies from patients in the RISK trial (GEO ID GSE57945)(iCD n=162, cCD n=56, UC n=62, control non-IBD group, n=42). **(b & c)** Following cell culture, T cells were washed and incubated with fluorescently labelled APC. Binding was measured by flow cytometry. Following APC pre-treatment, APC was removed, Th0 cells were washed with EDTA-containing PBS, and their capacity to initiate clotting was analysed by **(d-g)** thrombin generation and **(h**) clot formation assays. APC pre-treated Th1 cell-dependent thrombin generation was analysed by **(i-l)** thrombin generation assays and **(m)** clot formation assays. Following APC pre-treatment, T cell procoagulant activity was analysed by **(n)** FXa generation assay, **(o)** *F3* gene expression and **(p & q)** PDI cell surface expression by flow cytometry. Similarly, Th1 cell thrombogenicity was analysed by **(r)** FXa generation assay, **(s)** *F3* gene expression and **(t & u)** PDI cell surface expression by flow cytometry. Student-paired t-test, Student-unpaired t-test or Mann Whitney U Test was used where appropriate to determine statistical significance with *P≤0.05, **P≤0.01, ***P≤0.001 for 4-10 donors/group.

## DISCUSSION

IBD patients exhibit a significantly increased risk of VTE compared to the general population^4,5^, yet the molecular pathways underpinning the pathogenesis of VTE in IBD remain poorly understood. In this study, we report the presence of TF^+^ CD4^+^ T cells in the inflamed mucosa of paediatric IBD patients and colitogenic mice, uncover the thrombogenic phenotype of activated CD4^+^ T cells, and demonstrate a new role for APC signalling in mitigating CD4^+^ T cell thrombo-inflammatory activity.

Several studies have highlighted coagulation dysregulation in both IBD patients and pre-clinical models of IBD^12–14,26^. This disruption is particularly evident in models of innate immune-mediated colitis, characterised by enhanced procoagulant TF activity, microparticle release and diminished PC pathway activity in the gut epithelia^11,14,16^. This dysregulation was confirmed upon transcriptomic analysis of IBD patient colonic biopsy tissue, which revealed heightened TF expression in IBD patients compared to non-IBD patient control tissue. However, this bulk analysis did not provide insight into breadth of cellular sources of increased TF expression in IBD patients. Subsequent in silico analysis of innate and adaptive immune cell TF expression, however, suggested that adaptive immune CD4^+^ T cells may represent a surprisingly rich source of TF. CD4^+^ T cell TF expression is greater than that typically observed on more common cellular TF sources, such as activated monocytes or neutrophils^32,33^. As IBD pathophysiology is largely driven by T cell dysfunction^17^, we examined whether CD4^+^ T cells may represent a novel source of thrombogenic activity in IBD patients. We found that IBD patients exhibited significantly higher numbers of TF^+^ CD4^+^ T cells in their colons than non-IBD individuals. Furthermore, we observed increased TF expression on infiltrating immune cells and CD4^+^ T cells in the colons of mice in a T cell-mediated model of colitis. Consequently, TF^+^ CD4^+^ T cells exist in the gut and are significantly increased during active disease.

Cell surface TF is normally expressed in an ‘encrypted’ state, with limited procoagulant activity^23^. Several mechanisms that regulate TF adoption of a procoagulant, ‘decrypted’ state have been proposed. These steps typically either alter TF structural conformation (PDI-mediated disulphide bond rearrangement)^23^ or adjust the phospholipid microenvironment around TF to facilitate optimal tenase complex activity (PS and SM rearrangement)^23^. We observed that key mediators of TF decryption, namely ASMase membrane translocation, PDI surface expression and PS exposure, are activated in Th0 and Th1 cells, but largely absent in naïve T cells. Although stimuli for TF decryption in innate immune cells, such as TLR activation, cytokine stimulation or toxic stressors^32^, are unlikely to be shared with T cells, downstream signalling pathways leading to TF decryption are likely to overlap. For instance, activation of p38 mitogen-activated protein kinase (p38 MAPK)-dependent signalling in monocytes promotes TF decryption that is largely mediated by PS exposure^34^. Similarly, TCR engagement also promotes p38 MAPK signalling in CD4^+^ T cells^35^, which may also contribute to TF decryption. Interestingly, the expression of these decryption pathways have been implicated in several other aspects of T cell-mediated IBD pathophysiology. For example, ASMase expression in T cells enhances proliferation and survival following TCR activation^36^, specifically via co-stimulatory CD28 receptor ligation^37^. In addition, ASMase-deficient mice exhibit significantly higher numbers of splenic Tregs than wild-type mice^38^. Notably, pharmacological ASMase inhibition in DSS-induced colitis significantly reduces disease severity^39,40^. PS exposure on the T cell surface has been shown to bind T cell immunoglobulin and mucin domain containing-3 (TIM-3) and act as a co-receptor for T cell activation^41^. Furthermore, PS externalized by antigen-stimulated CD8 T cells is linked to T cell activation, with increased expression of CD69 and levels of IFN-γ, IL-2, and TNF-α reported^42^. In trinitrobenzene sulfonic acid (TNBS)-induced colitis, PS inhibition by annexin A5 alleviated disease via reduced inflammatory cell infiltration due to inhibition of endothelial cell activation^43^. Furthermore, pups reared on artificial milk with the addition of lactadherin, the PS binding and blocking glycoprotein, displayed increased levels of tolerogenic CD3^+^CD4^+^CD25^+^ T cells in their Payer’s patches compared to pups reared on artificial milk alone^44^. There are also several reports indicating a role for PDI in regulating T cell responses in cancer and infection. PDIA3-specific T-cell clones were found in colorectal cancer (CRC) patients and displayed aberrant immunity in models of malignant melanoma^45^. Furthermore, recent reports have suggested an important role for PDI in eliciting enhanced anti-parasitic responses in T cells. Immunization with *Leishmania donovani* PDI (*Ld-*PDI) followed by challenge with the *L. donovani* resulted in enhanced CD4^+^ Th1 and Th17, and CD8^+^ T cell immunoreactivity via MAPK-pathway signalling in Balb/c mice^46^. Similarly, in studies of *Toxoplasma gondii* infection, Balb/c mice immunized with recombinant *T. gondii* PDI (r*Tg-*PDI) showed enhanced Th1 immunity and reduced levels of parasitic infection following challenge with *T. gondii*^47^. Although our study used *ex vivo*-generated Th0 and Th1 cells to assess T cell procoagulant activity and TF decryption, the specific inflammatory and cellular determinants that drive activation of T cell thrombogenicity *in vivo* remain unknown and represent an important avenue for further investigation.

APC is best characterised as a plasma anticoagulant that degrades activated cofactors factor V (FVa) and factor VIII (FVIIIa) to limit thrombin generation and subsequent fibrin deposition^24^. Like other coagulation proteases, APC mediates receptor-mediated cell signalling that is predominantly anti-inflammatory^48^. APC signalling is typically mediated by activation of protease-activated receptors (PARs), in particular PAR1 and PAR3. PAR signalling by APC typically requires APC co-localisation via binding to a co-receptor, most commonly EPCR^49^. Notably, disruption of APC generation or activity has been shown to contribute to T cell-mediated inflammatory diseases^28–30^. APC inhibits pro-inflammatory Th17 and Th1 activity and promotes the expansion of tolerogenic Tregs^25–28,50^. APC administration ameliorates graft vs host disease (GvHD) via anti-inflammatory PAR1 signalling on T cells^50^. Furthermore, pre-incubation of donor T cells with APC *ex vivo* inhibits allogenic T cell expansion and increases the pool of Tregs before transplantation via PAR2 and PAR3 signalling^28^. AP also promotes altered T cell metabolism, resulting in *Foxp3* promoter demethylation^30^. Furthermore, EPCR-deficient (*PROCR*^low^) mice (that generate less APC and have limited capacity to facilitate EPCR-dependent APC signalling) exhibit exacerbated disease in experimental autoimmune encephalomyelitis, via enhanced pro-inflammatory Th17 cell generation^29^. Similarly, in preclinical models of atopic dermatitis and arthritis, APC administration reduces Th1 and Th17 cell populations via PAR1 and PAR2 signalling^25–27^. Our data suggest that, in addition to these potent anti-inflammatory activities, diminished APC activity may contribute to aberrant mucosal haemostasis and impair local regulation of T cell-mediated thrombogenicity.

In conclusion, we show for the first time that APC binding to the surface of CD4^+^ T cells and subsequent signalling activity directly regulates CD4^+^ T cell-expressed TF procoagulant activity. The ability of APC to limit disease-associated features of IBD in preclinical colitis models has led to its consideration as a therapy in treating IBD^14,16^. However, the potent anticoagulant activity of APC and its association with increased bleeding risk in patients represent a significant obstacle to its successful clinical implementation. Our study, however, suggests that recombinant ‘non-anticoagulant’ APC variants that are unable to degrade FVa and FVIIIa effectively may still selectively regulate T cell-specific immunothrombotic activity, without impacting canonical APC anticoagulant substrates. Consequently, these findings highlight a potential therapeutic role for APC targeting CD4^+^ T cell thrombo-inflammatory activity in IBD and other T cell-mediated disease contexts.

## ACKNOWLEDGEMENTS

Grant support for RJSP is provided by Science Foundation Ireland (21/FFP-A/8859), The National Children’s Research Centre (C/18/3) and Health Research Board (ILPPOR-2022-060).

## AUTHOR CONTRIBUTIONS

G.L. and R.J.S.P. devised the study and experimental strategy and wrote the manuscript. G.L., P.A.K., A.M.R., S.H., S.C.B., J.S.O’D, and P.T.W. performed experiments and analysed data. All authors reviewed and contributed to the final version of the manuscript.

## SUPPLEMENTARY FIGURE LEGENDS

**Supplementary Figure 1:**
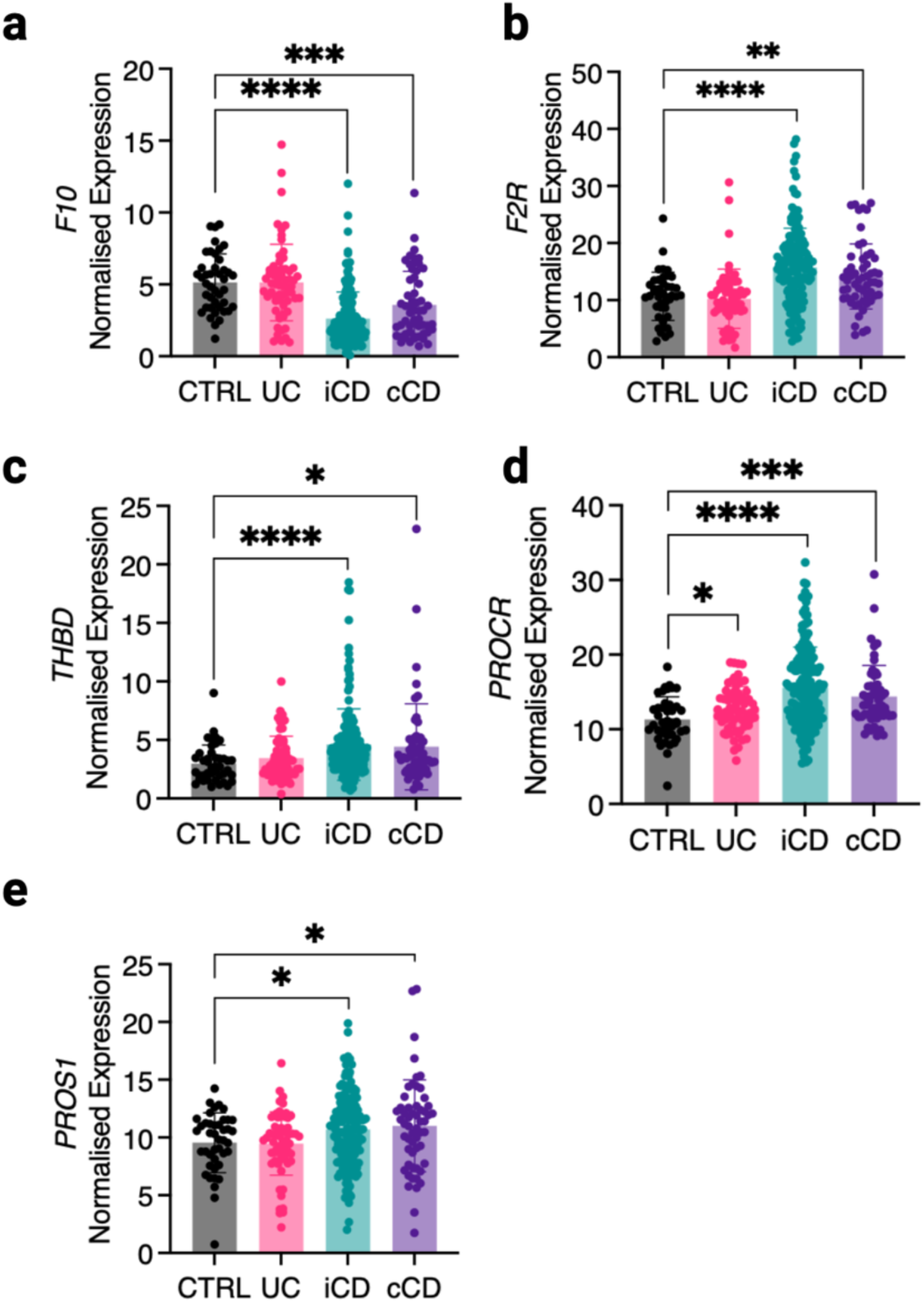
Coagulation genes are dysregulated in IBD. Expression of **(a)** *FX*, **(b)** *F2R*, **(c)** *THBD*, **(d)** *PROCR* and **(e)** *PROS1* from RNA-seq data of colonic biopsies from Crohns Disease (CD; illeal (iCD, n=162), colonic (cCD, n=56), Ulcerative Colitis (UC, n=62), and query IBD healthy control cohort (CTRL, n=42) patients in the RISK trial (GEO ID GSE57945). Student t-test or Mann Whitney U Test was used where appropriate to determine statistical significance with *P≤0.05, **P≤0.01, ***P≤0.001.

**Supplementary Figure 2:**
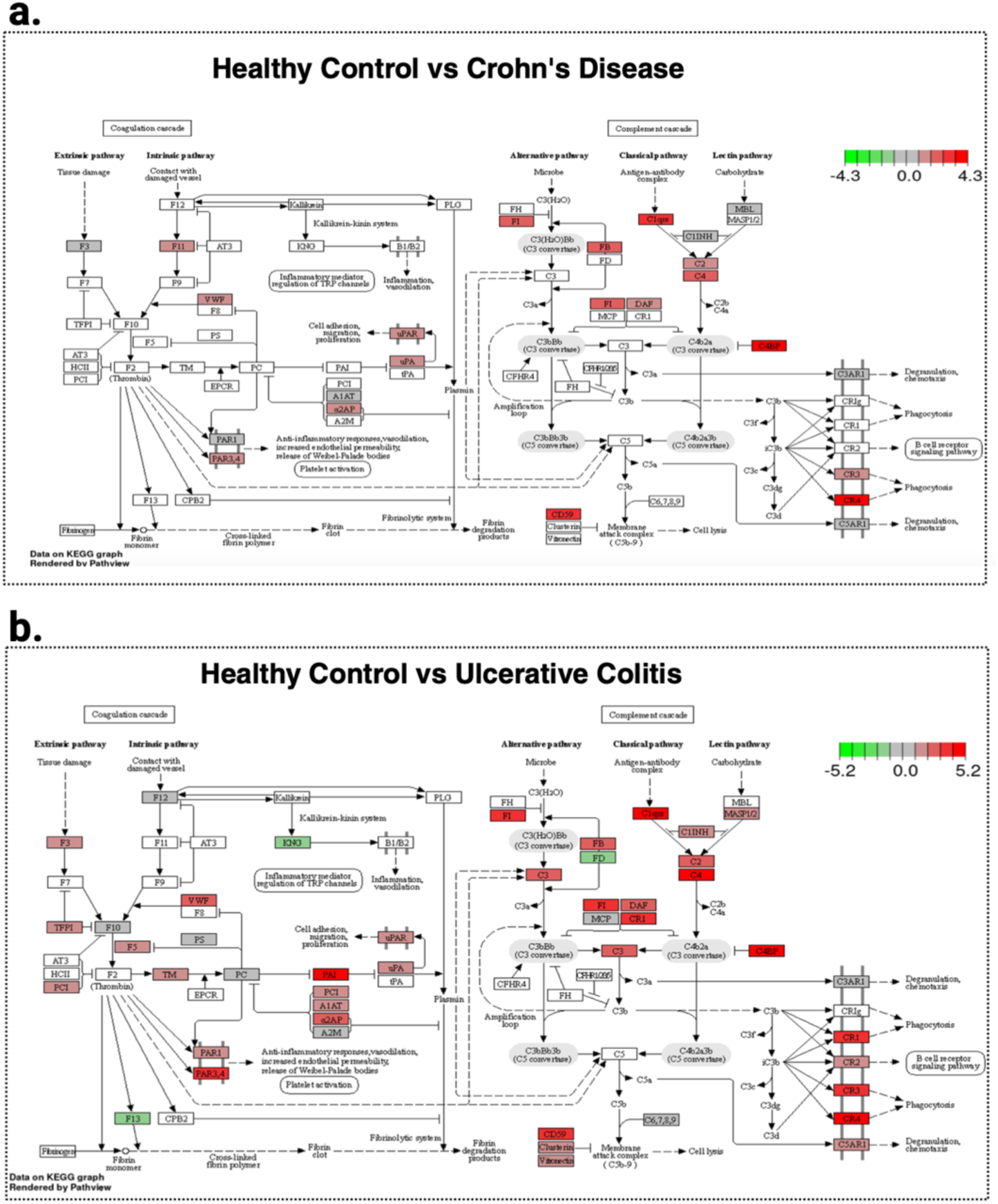
Coagulation pathways are dysregulated in IBD patients’ colons. KEGG pathway analysis was used to depict dysregulation in the coagulation pathway in RNA-seq samples of colonic biopsies from paediatric CD patients compared to **(a)** healthy controls and **(b)** paediatric UC patients CD, n=11; UC, n = 7; Ctrl, n = 12.

**Supplementary Figure 3:**
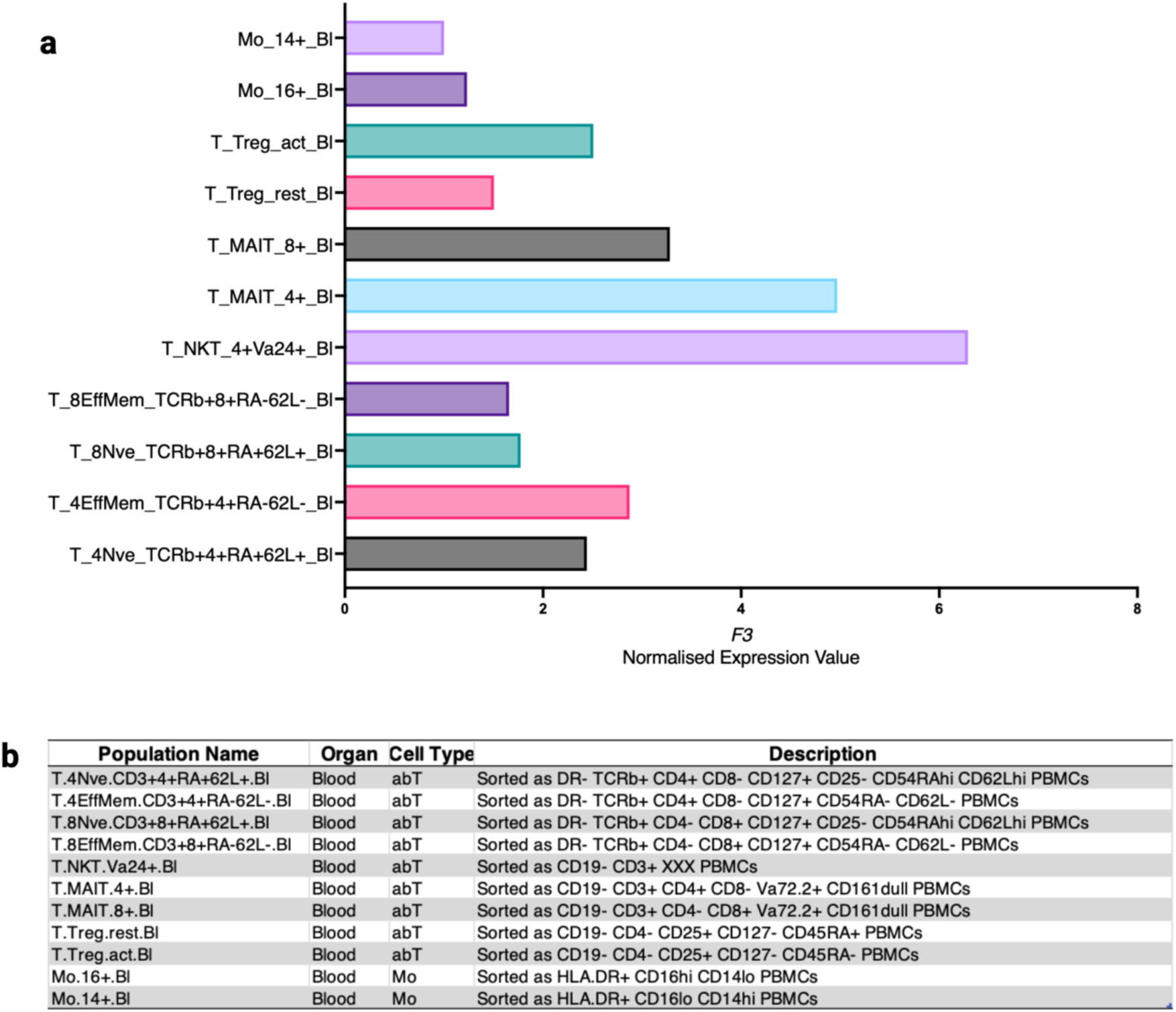
*F3* expression by human immune cells. **(a)** TF expression in human immune cells contributing to IBD pathogenesis was analysed using the Human Cell Atlas of RNA-seq datasets uploaded to the Immunological Genome Project (http://immunecellatlas.net). **(b)** A descriptive table of immune cell populations and their markers was included in the analyses.

**Supplementary Figure 4:**
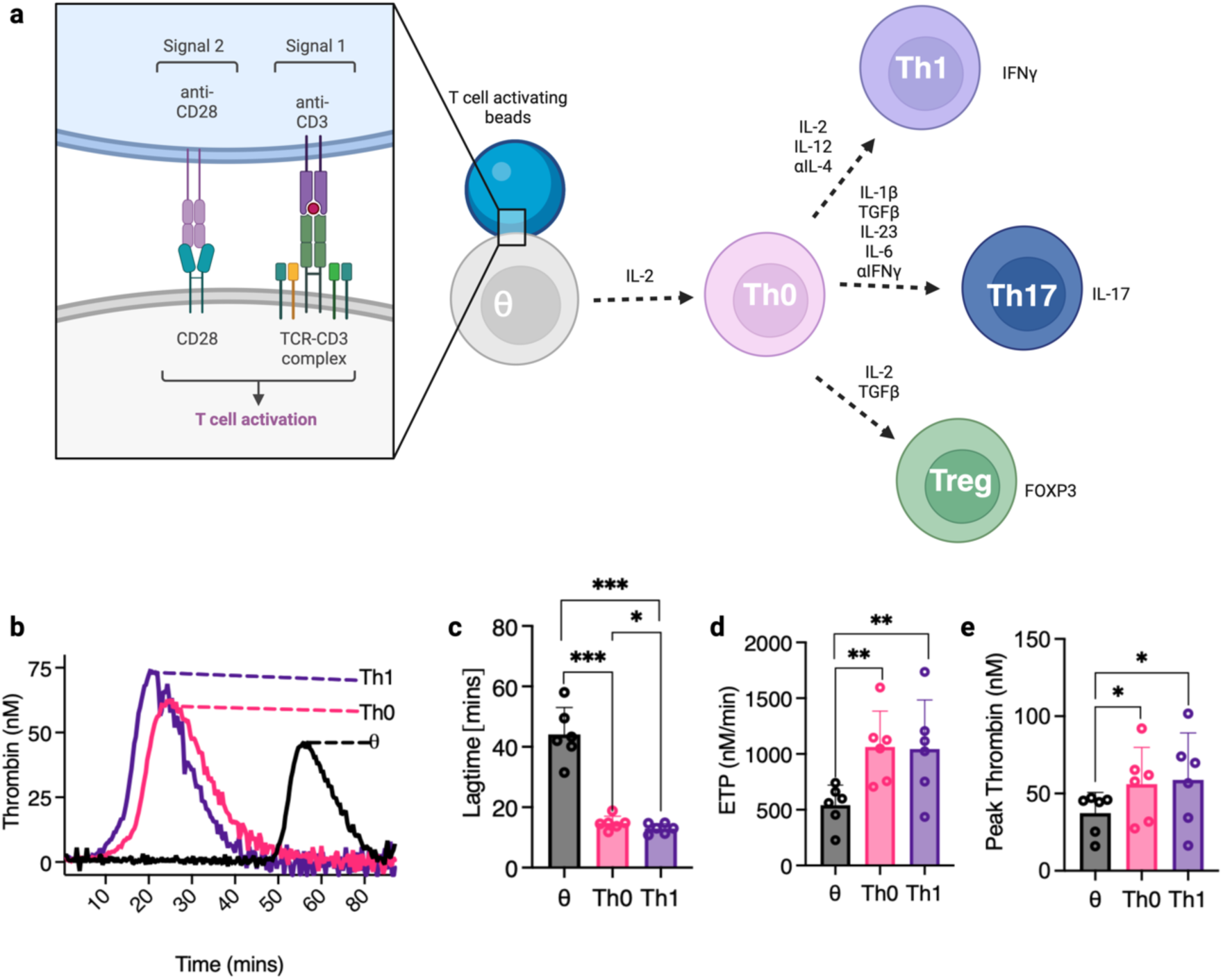
CD4^+^ T cell activation, differentiation, and thrombin generation in FXII-deficient plasma. **(a)** Schematic diagram representing the process of T cell activation and differentiation requirements. CD4^+^ T cells require 2 signals to activate fully. They first require stimulation of the T cell receptor (TCR/CD3)(signal one) by Major histocompatibility complex (MHC) Class 2 molecules presented by antigen-presenting cells or by anti-CD3 molecules presented on the surface of synthetic beads. Stimulation of the T cell co-stimulatory receptor CD28 is then required (signal 2), either by CD86 presented by antigen-presenting cells or by anti-CD28 molecules presented on the surface of synthetic beads. In our experiments, we used anti-CD3/anti-CD28 coated beads to activate T cells fully. Once activated, IL-2 is required to enhance viability and proliferation, and specific cytokines are applied to skew the activated Th0 cells towards distinct inflammatory lineages (Th1: IL-2, αIL-4, IL-12, Treg: IL-2, TGFβ, Th17: TGFβ, IL-1β, IL-23, & IL-6, αIFNγ). **(b-e)** Calibrated automated thrombinography was used to measure extrinsic pathway-mediated T cell procoagulant activity using FXII-deficient plasma. **(c)** Lagtime, **(d)** endogenous thrombin potential (ETP) and **(e)** peak thrombin levels were measured and compared between θ, Th0 and Th1 cells. Student-paired t-test was used to determine statistical significance with *P≤0.05, **P≤0.01, ***P≤0.001 for 6 donors/group.

**Supplementary Figure 5:**
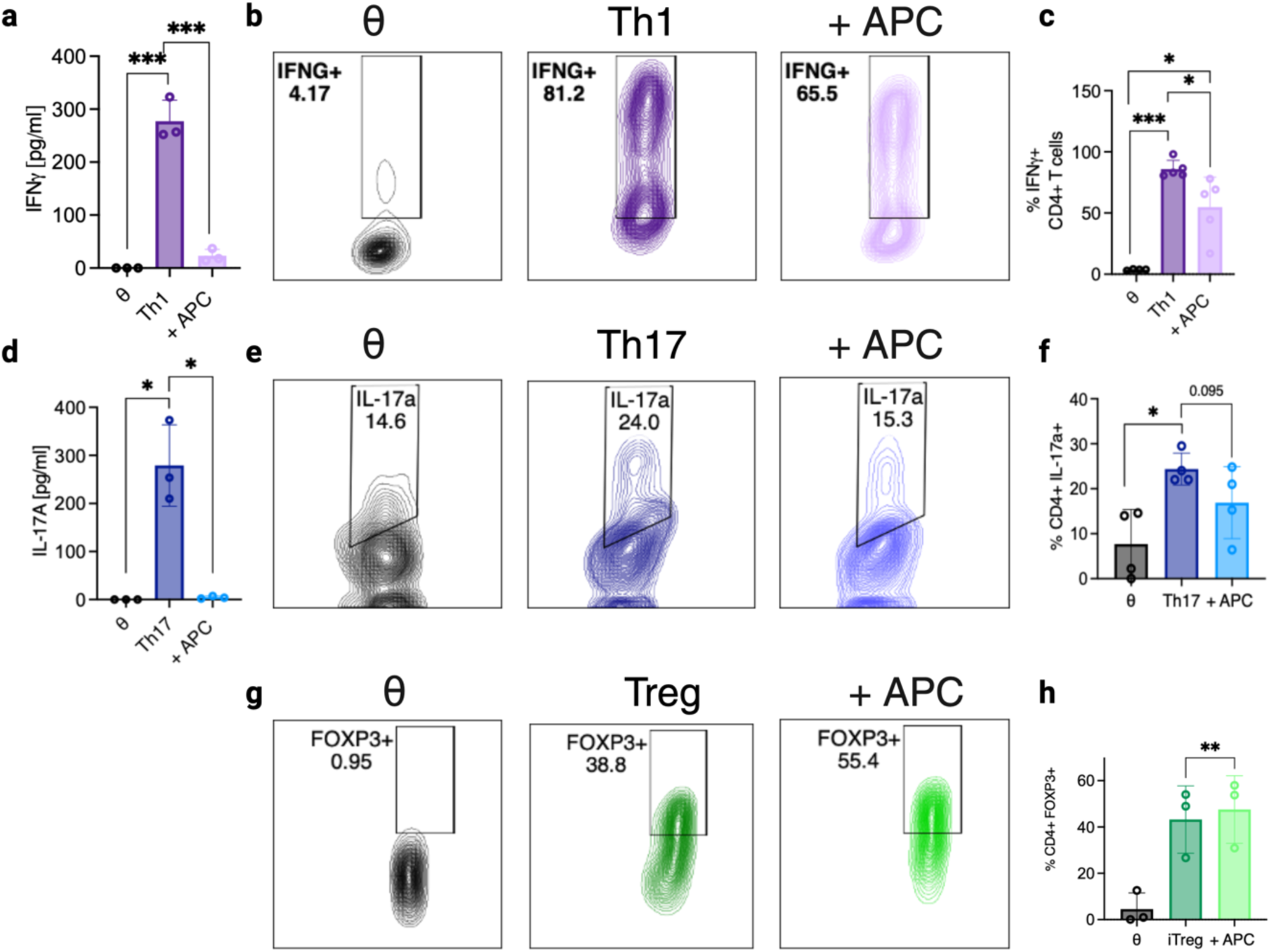
APC regulates CD4^+^ T proinflammatory activity. **(a-h)** CD4^+^ T cells were isolated from donor human blood and plated at a density of 0.8x10^6^/ml with IL-2 for unactivated conditions (θ). Plated cells were activated with the addition of αCD3/αCD28 activation beads and stimulated with IL-2 for Th0 conditions, differentiation cytokines were added to skew cells to a **(a-c)** Th1 lineage (αIL-4 + IL-12, a **(d-f)** Th17 lineage (TGFβ, IL-1β, IL-23, & IL-6, αIFNγ) or a **(g & h),**Treg lineage (IL-2, TGFβ), and cells were treated +/-APC for 5 days. In Th1 cells pre-treated with APC (**a)** IFNγ levels were measured by ELISA, and **(b & c)** the percentage of IFNγ^+^ cells was measured by flow cytometry. In Th17 cells that were pre-treated with APC **(d)** IL-17a levels were measured by ELISA, and **(e & f)** the percentage of IL-17a^+^ cells was measured by flow cytometry. The percentage of FOXP3^+^ cells was measured by flow cytometry in Treg cells that had been previously treated with APC **(g & h)**. Student-paired t-test was used to determine statistical significance with *P≤0.05, **P≤0.01, ***P≤0.001 for 3-5 donors/group.

## SUPPLEMENTARY MATERIALS AND METHODS

### ELISAs

An ELISA kit for human IFNγ was purchased from eBioscience (Thermofisher) and performed according to the manufacturer’s instructions using Corning High Binding ELISA plates (Merck). All ELISAs were analysed using a Synergy MX microplate reader (BioTek).

### mRNA isolation and RT-qPCR of CD4^+^ T cells

Following T cell culture, supernatants were removed, and cells were washed once with PBS and transferred to a 1.5ml RNAse free Microfuge Tubes (Invitrogen). Cells were then lysed using 350µL lysis buffer, and total RNA was isolated according to the manufacturer’s instructions (Ambion PureLink RNA isolation kit, Thermofisher). Gene expression was determined by RT-qPCR in duplicate with Power Up SYBR green master mix (Thermofisher) using a 7500 Fast system (Applied Biosystems). Relative-fold changes in mRNA expression were calculated using the cycle threshold (C_T_) and normalised to the *RPS18* housekeeping gene. Primer sequences are listed in **Supplementary Table 1**.

### Flow cytometry

All fluorescently-labelled antibodies were purchased from Thermofisher (αCD4, αIFNg) and Biolegend (αEPCR). Cell viability was measured using the LIVE/DEAD Fixable Dead Cell Stain Kit (Aqua, Scarlet & Near IR) (Invitrogen, Thermofisher). Before staining, cells were incubated with an anti-CD16/CD32 monoclonal antibody to block Fc receptors (Invitrogen). Cells were washed using PBS and incubated with LIVE/DEAD dye for 30 minutes at 4°C. Cells were washed using PBS supplemented with 2% FBS and then incubated with fluorescently-labelled antibodies for 1 hour at 4°C. For unconjugated antibodies (αASM (Invitrogen) & αTF (R&D)), cells were incubated for 1 hour at room temperature with the primary antibody, washed and incubated with a specific secondary antibody for 1 hour at 4°C (Donkey anti-Goat IgG (H+L) Cross-Adsorbed Secondary Antibody Alexa Fluor 555 and Invitrogen Goat anti-Rabbit IgG (H+L) Highly Cross-Adsorbed Secondary Antibody Alexa Fluor 555, Thermofisher). Fluorescence Minus One (FMO) and/or isotype controls were used to assess positive staining. Intracellular protein expression was evaluated by first restimulating the cells with phorbol-12-myristate 13-acetate (PMA; 10 ng/ml) (Sigma Aldrich), Ionomycin (1 ug/ml) (Sigma Aldrich) and Brefeldin A (5 ug/ml) (eBioscience) for 4–6 h at 37 °C. A FOXP3 staining buffer set (eBiosciences) was used in accordance with the manufacturer’s instructions to fix and permeabilise cells after surface staining to facilitate the detection of intracellular cytokines. Intracellular fluorescently labelled antibodies were incubated for 1 hour at 4°C. Multi-parameter analysis was then performed on an LSR/Fortessa (BD) or Attune Nxt (Thermofisher) and analysed using FlowJo™ 10 software. Antibodies and dyes are listed in **Supplementary Table 2**.

### Cell binding assays

Recombinant human lactadherin (R&D Systems) and human APC (Cambridge ProteinWorks) were fluorescently labelled using Lightning-Link® Rapid Alexa Fluor 488 Antibody Labelling Kits in accordance with the manufacturer’s instructions (Novus Biologicals, R&D). Before staining, cells were first incubated with an anti-CD16/CD32 Monoclonal Antibody to block FC receptors (Invitrogen™, Thermofisher). Cells were washed using PBS and incubated with LIVE/DEAD™ dye for 30 minutes at 4°C. Cells were washed using PBS supplemented with 2% FBS and incubated with CD4 antibodies for 30 minutes at 4°C. Cells were washed and incubated with the fluorescently labelled proteins for 2 hours. Fluorescence Minus One (FMO) and/or isoclonic controls were used to assess positive staining. Multi-parameter analysis was then performed on an LSR/Fortessa (BD) or Attune Nxt (Thermofisher) and analysed using FlowJo™ 10 software.

### Immunofluorescence

Paraffin-embedded blocks of colon biopsies from children diagnosed with CD (n =2), UC (n =2) or healthy controls (n =3) were obtained from Our Lady’s Children’s Hospital Crumlin. Blocks were sectioned to 5 μm thickness using a microtome and mounted on Superfrost Plus adhesion slides (Thermofisher). Antigen retrieval was performed using IHC Antigen Retrieval Solution (eBioscience, Thermofisher) in a microwave. Tissue was probed with α-human CD3 (FITC)(10ug/ml) (eBioscience, Thermofisher), α-human CD4 (PE)(10ug/ml) (eBioscience, Thermofisher), α-human TF (2.5ug/ml) (R&D), or isotype controls. Paraffin-embedded blocks of colons from *Rag1*^−/−^ mice adoptively transferred with wild-type T effector cells, or PBS, were sectioned to 5 μm thickness using a microtome and mounted on Superfrost Plus adhesion slides (Thermofisher). Antigen retrieval was performed using 10 mM citrate buffer in a microwave. Tissue was probed with α-mouse CD3 (1:100)(FITC) (eBioscience, Thermofisher), α-mouse TF (2.5ug/ml) (R&D) or isotype controls. The secondary antibodies used were donkey anti-Goat IgG (H+L) Cross-Adsorbed Secondary Antibody Alexa Fluor™ 555 (1ug/ml) and Invitrogen Goat anti-Rabbit IgG (H+L) Highly Cross-Adsorbed Secondary Antibody Alexa Fluor™ 555 (1ug/ml). Tissue was mounted using SlowFade Gold antifade Mountant with DAPI (Thermofisher). Images were taken using a confocal microscope Zeiss LSM700. Positive cells stained with αCD3, αCD4 and αTF were quantified using the ImageJ tool Fiji and the Count plugin. An average of three different images per sample were analysed statistically using GraphPad Prism 9.5 software to give an average number of cells/field. Antibodies and dyes are listed in **Supplementary Table 2**.

### RNAseq data analysis

We utilised multi-omic data from publicly available datasets of IBD patient intestinal biopsies, including RNA-seq data from the Risk Stratification and Identification of Immunogenetic and Microbial Markers of Rapid Disease Progression in Children with Crohn’s Disease (RISK) study, in which we compared the expression of genes of interest between paediatric CD and UC patients and healthy controls . We accessed NCBI Gene Expression Omnibus datasets using GEO IDs GSE57945 (Risk Cohort). The log2 fold-change and p-value significance data were downloaded and analysed using GraphPad Prism 9.5 software. Data are shown as means ± SEM.

We also utilised in-house RNA-seq data generated from paediatric IBD patients and control participants’ biopsies recruited in the DOCHAS study at the Children’s Health Ireland (CHI) gastroenterology unit. Data was analysed in R. Relative gene expression per individual gene is shown as z scores in a heatmap. KEGG pathway analysis was used to depict dysregulation in the coagulation pathway.

---

**SUPPLEMENTARY TABLE 1:**
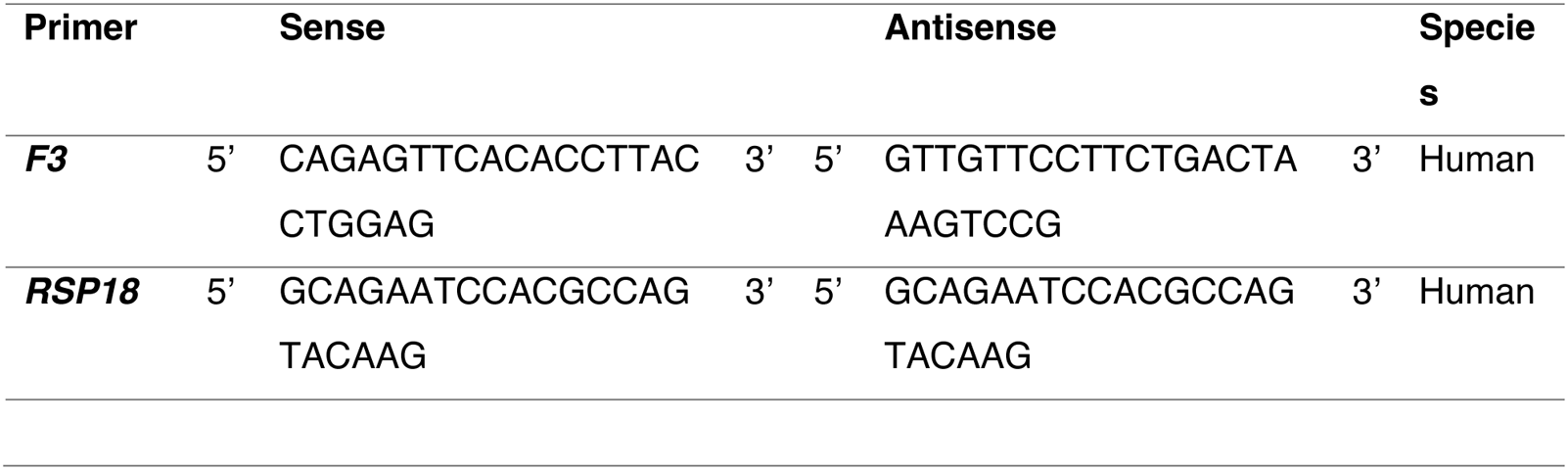

**SUPPLEMENTARY TABLE 2:**
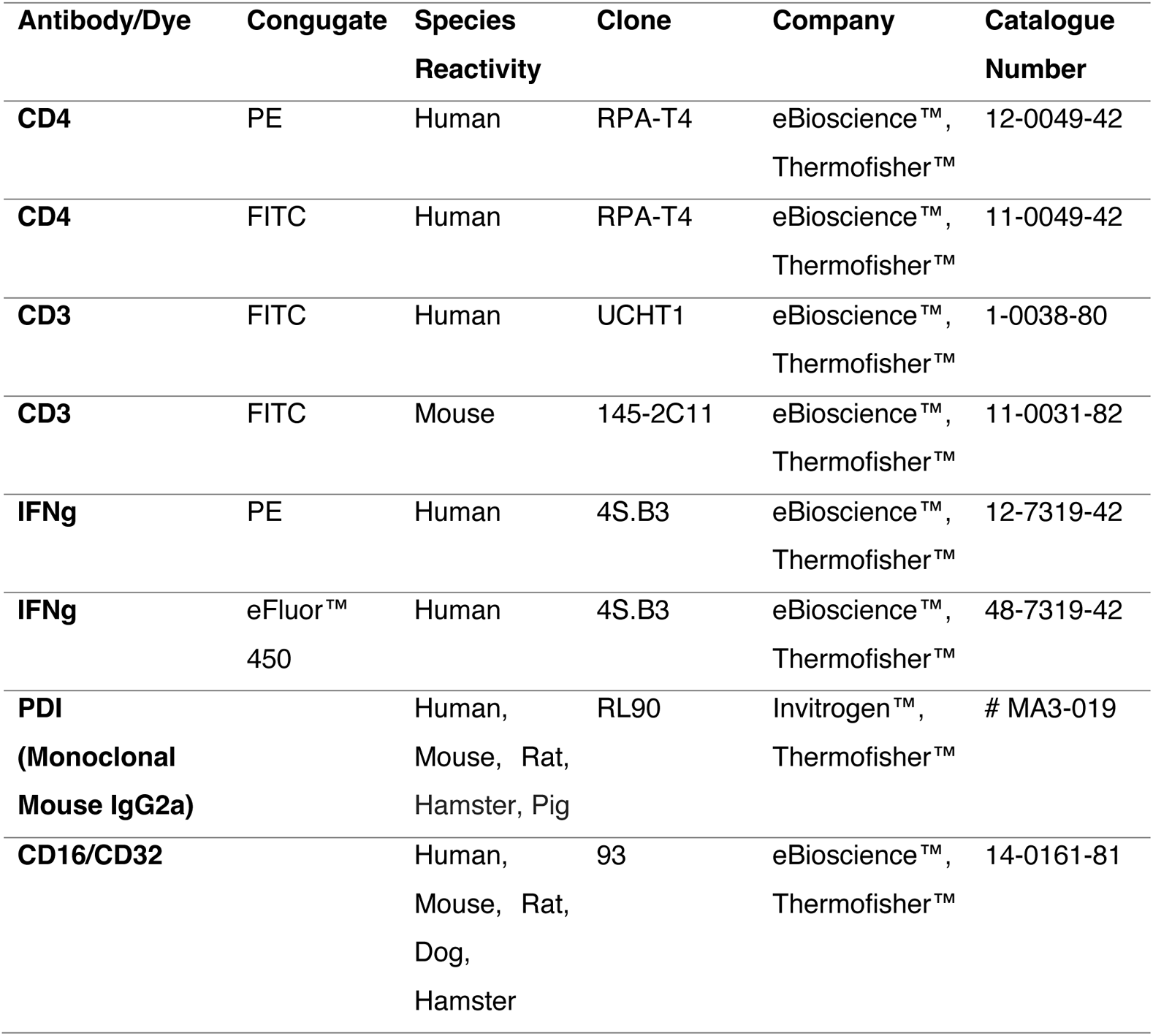
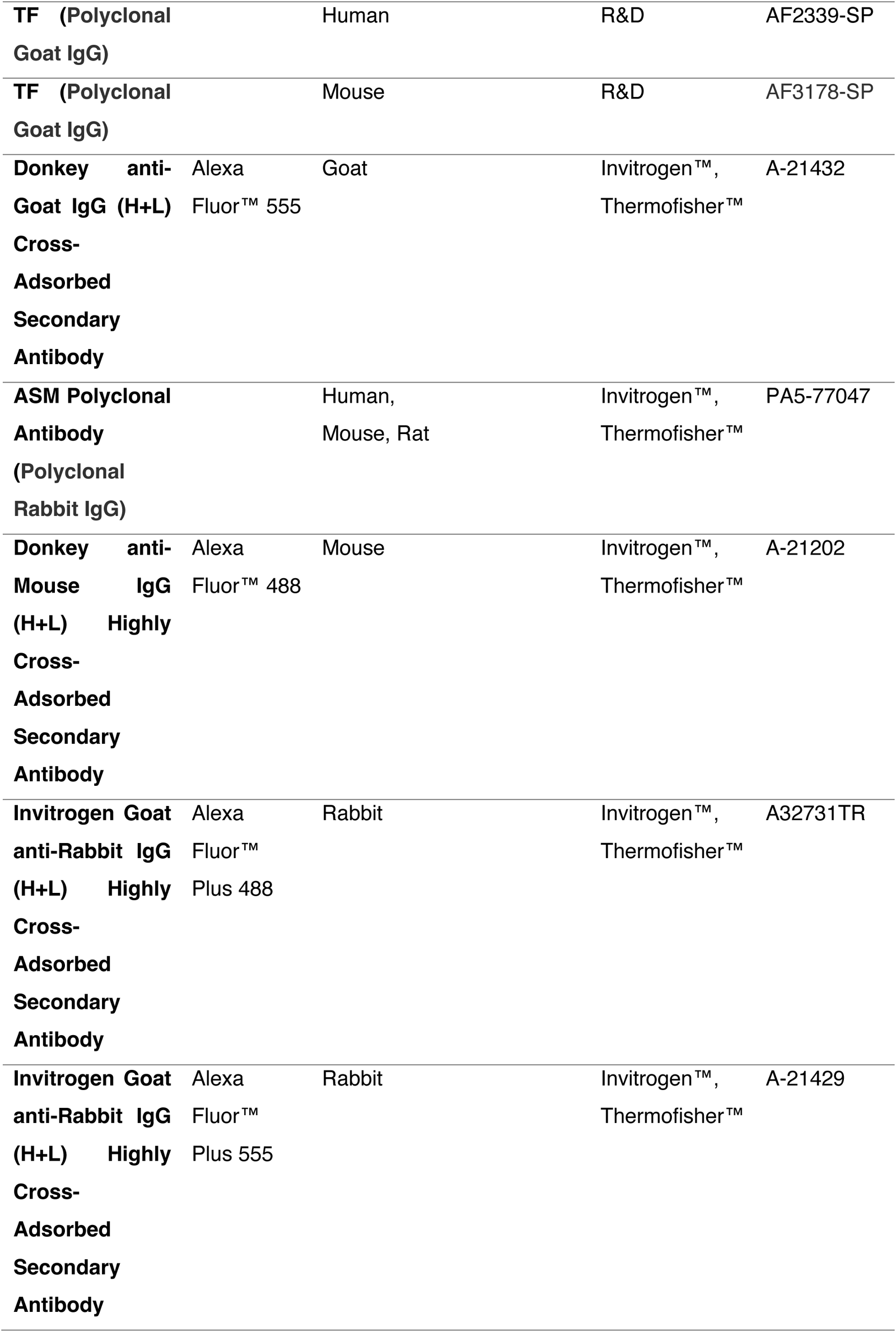
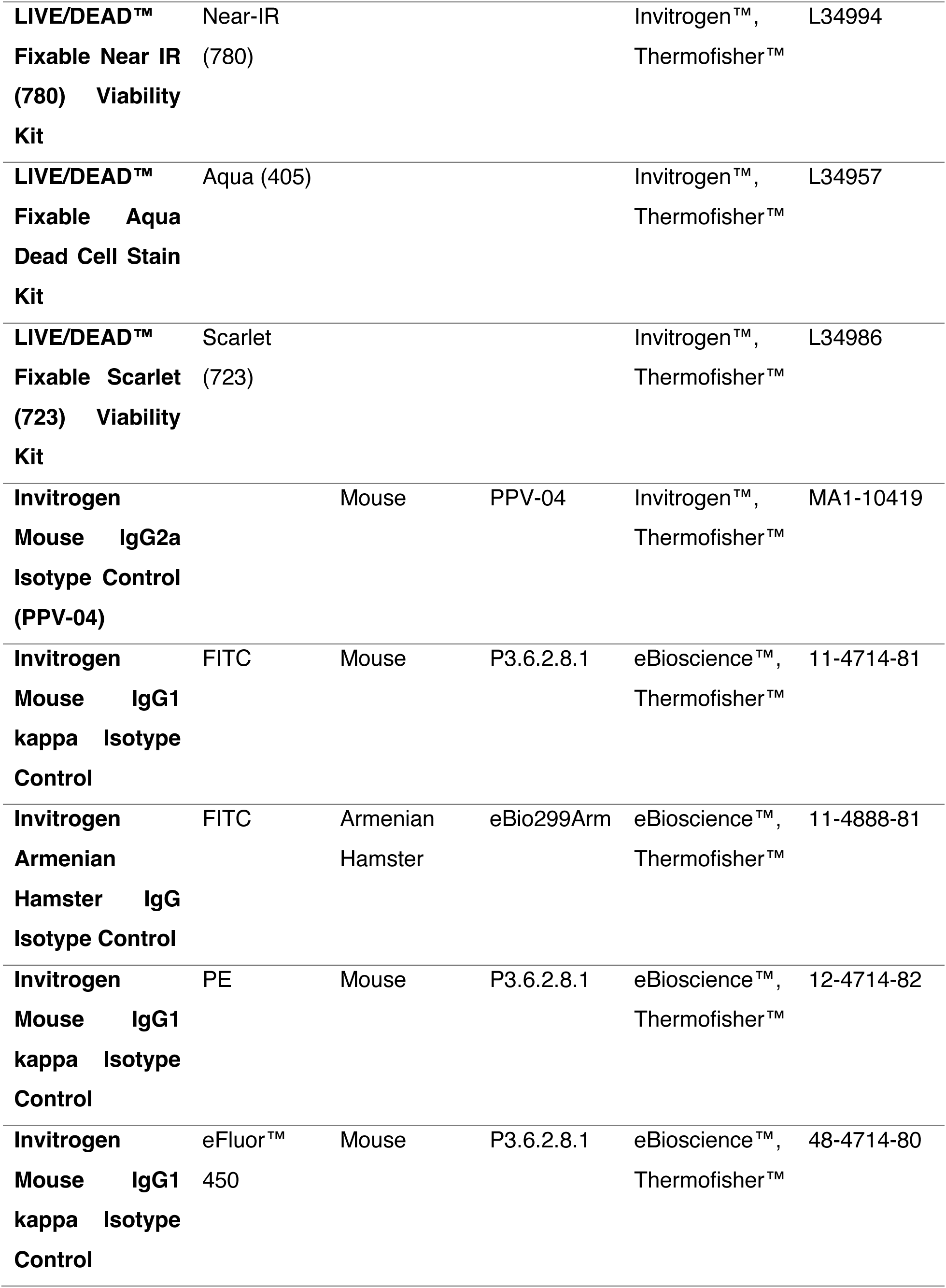
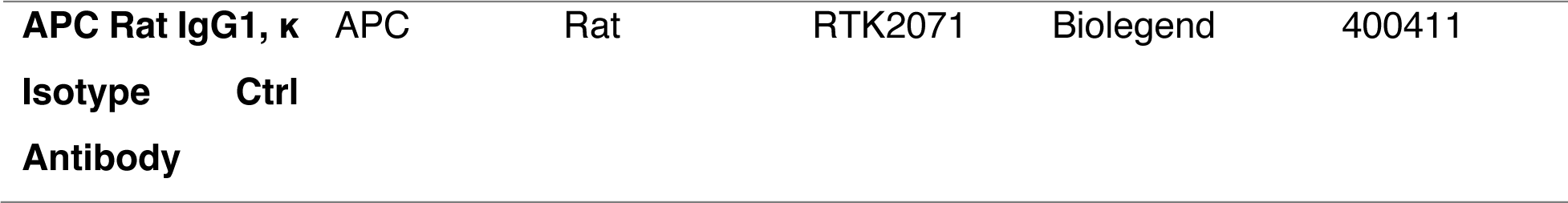

